# Machine learning predicts lifespan and underlying causes of death in aging *C. elegans*

**DOI:** 10.1101/2024.03.20.585803

**Authors:** Carina C. Kern, Petru Manescu, Matt Cuffaro, Catherine Au, Aihan Zhang, Hongyuan Wang, Ann F. Gilliat, Marina Ezcurra, David Gems

**Affiliations:** Institute of Healthy Ageing, and Research Department of Genetics, Evolution and Environment, University College London, London UK; Department of Computer Science, Faculty of Engineering Sciences, University College London, London UK; School of Biosciences, Stacey Building, University of Kent, Canterbury, Kent, UK

**Keywords:** Age-related disease, aging, *C. elegans*, lifespan, machine learning, reproductive death

## Abstract

Senescence (aging) leads to senescent pathology that causes death, and genes control aging by determining such pathology. Here we investigate how senescent pathology mediates the effect of genotype on lifespan in *C. elegans* by means of a data-driven approach, using machine learning (ML). To achieve this we gathered extensive data on how diverse determinants of lifespan (sex, nutrition, genotype) affect patterns of age-related pathology. Our findings show that different life-extending treatments result in distinct patterns of suppression of senescent pathology. By analysing the differential effects on pathology and lifespan, our ML models were able to predict >70% of lifespan variation. Extent of pathology in the pharynx and intestine were the most important predictors of lifespan, arguing that elderly *C. elegans* die in part due to late-life disease in these organs. Notably, the mid-life pathogenetic burst characteristic of hermaphrodite senescence is absent from males.

## Introduction

Human senescence is largely studied from the perspective of age-related pathologies and diseases, but less so in terms of the primary causes of aging. Conversely, the genetics of aging in *C. elegans* has identified signalling pathways with effects on lifespan that show evolutionary conservation^1^, but the pathologies involved remain poorly understood. The large differences in maximum lifespan between animal species^2, 3^ demonstrates that the aging process is to a large extent genetically determined. A major objective of *C. elegans* aging studies is to establish how gene function determines senescence, and thereby arrive at an understanding of its mechanistic basis, including the causes of late-life disease. In our current knowledge of the chain of events in *C. elegans* leading from gene action to lifespan, the most poorly understood links involve the causes (etiologies) of senescent pathologies (including late-life diseases), and how late-life pathology causes death (Figure 1a).

**Figure 1.**
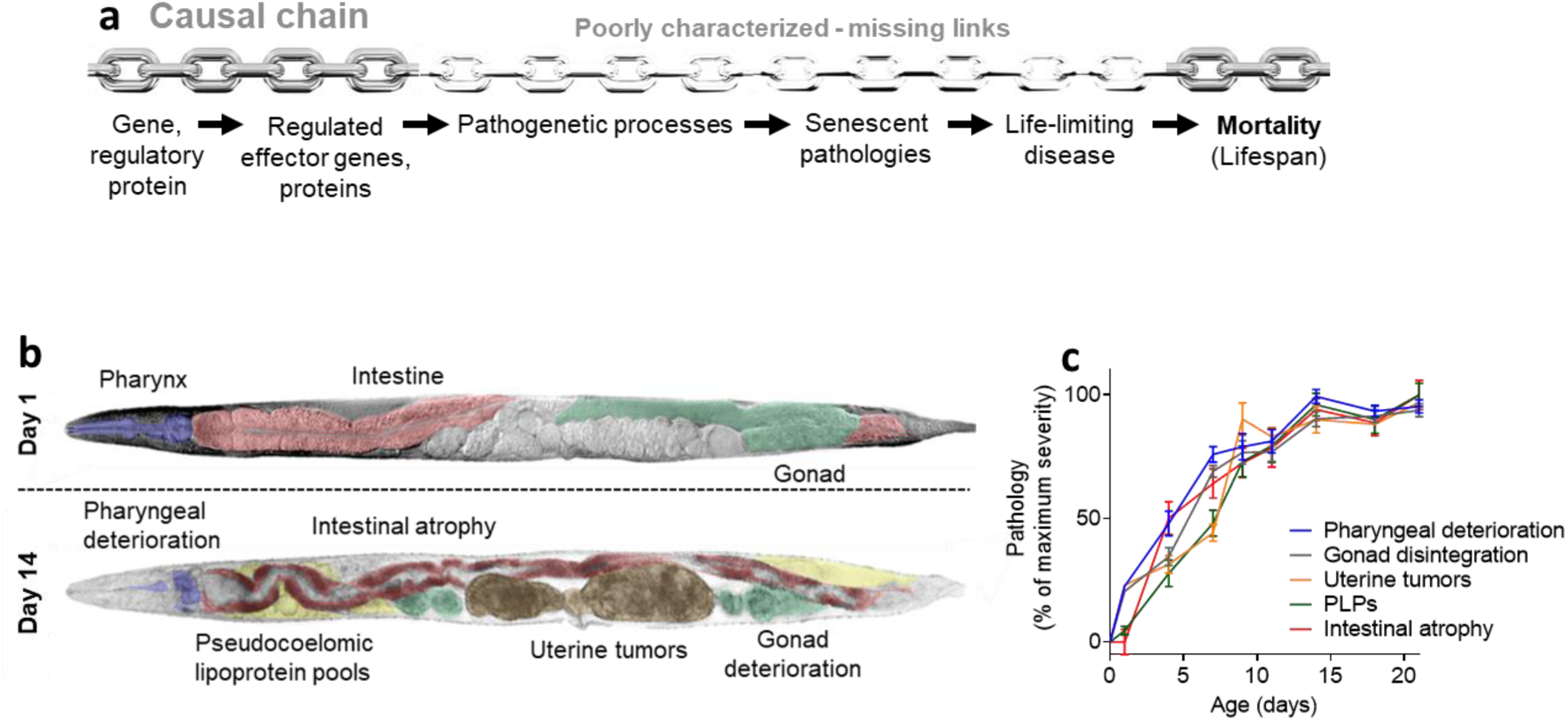
Senescent pathogenesis is a missing link in the chain of events between gene action and lifespan. (a) The causal chain of events between gene action, aging and lifespan. Senescent disease etiology is typically multifactorial and influenced by multiple factors^4^. (b) Pathological changes from day 1 to day 14 visibly affect the pharynx, intestine, gonad, and uterus (uterine tumors) and cause ectopic deposition of lipid/yolk in the body cavity (pseudocoelomic lipoprotein pools, PLPs). DIC images of worms with artificial coloring of organs/pathological changes. Top: Young, healthy animal with intact tissues and no pathology. Bottom: Old animal with multiple pathologies. (c) A burst of pathogenesis in early-mid adulthood. Development of age-related pathologies in *C. elegans* hermaphrodites starts in early adulthood (day 1-4), largely attaining their full extent by ∼day 10^5^.

Some knowledge of *C. elegans* senescent pathology exists thanks to a small number of pioneering studies using techniques such as light (especially Nomarski) microscopy, automated image analysis and electron microscopy^6, 7, 8, 9, 10, 11^. However, senescent pathology has been little quantified during aging in long-lived (Age) mutants, or understood in terms of its underlying mechanisms (etiologies).

We have taken a systematic approach, using Nomarski microscopy, to measure the development of naturally-occurring pathologies during nematode aging, and to survey effects of a range of interventions which alter lifespan. Using a similar but more piecemeal and less extensive approach we previously identified a putative cause of intestinal atrophy and pseudocoelomic lipoprotein pools (PLPs, or yolky pools): autophagy-assisted repurposing of intestinal biomass to generate yolk^5, 12, 13^.

Our main methodology builds upon earlier work^6, 7, 9, 14, 15, 16^ that has identified and characterized five prominent senescent pathologies that are observable in aging *C. elegans*: deterioration of the pharynx, atrophy of the intestine, PLP accumulation, atrophy and fragmentation of the distal gonad, and development of teratoma-like uterine tumors (Figure 1b,c, Supplementary Figure 1a-c, Supplementary Table 1; see Methods for details of scoring)^5, 7, 16, 17, 18^. *C. elegans* cohorts are aged under standard laboratory conditions (with *Escherichia coli* OP50 as a food source, at 20°C), and imaged at intervals from early adulthood onwards until all have died of old age.

Typically during animal aging (as in our own species) age-related diseases appear increasingly towards the end of life, leading to death. By contrast, wild-type *C. elegans* hermaphrodites exhibit a burst of pathogenesis in early-mid adulthood, driven by gene function^5, 19, 20, 21, 22^. A recently proposed hypothesis with some evidential support is that this reflects a semelparous life history in *C. elegans* hermaphrodites. In this, a suicidal reproductive effort (including self-destructive biomass repurposing to generate yolk that is vented to support larval growth) leads to reproductive death^18, 23, 24, 25^.

In this study we describe this pathogenetic burst in more detail, and examine how it is affected by sex and diverse treatments that alter lifespan, including altered insulin/IGF-1 signaling, protein synthesis and mitochondrial status. Artificial intelligence(AI)/ machine learning (ML) methods are proving increasingly powerful at yielding biological insight, as exemplified by the capacity of AlphaFold to predict protein function from primary structure^26^. In research on aging in *C. elegans*, ML has begun to be applied to the prediction of drugs that extend lifespan^27, 28^. Here we take the novel approach of applying AI/ML data-driven approaches to study the mechanistic basis of aging in *C. elegans*, specifically to explore the relationship between senescent pathology and lifespan across a range of genotypes and conditions.

## Results

### Life-extending treatments differentially affect pathologies of aging

For this ML analysis of the relationship between senescent pathology and lifespan in *C. elegans* we first gathered and compiled data. This included previously published data, and data newly generated for the purpose of this study. For a full listing of all data and sources see Supplementary Table 2. The data comprised measurements of severity of pathologies at multiple time points during adulthood, and lifespan estimates. In most cases, these were mean values for population cohorts, but data from individual wild-type *C. elegans* was also included. Population cohort studies included diverse mutations and RNAi treatments in *C. elegans*. Also included was data from different species in the *Caenorhabditis* and *Pristionchus* genera; notably, hermaphrodites show far higher levels of pathology and are shorter lived than females of sibling species^18^.

New data gathered for this study included the following treatments. Culture on axenic medium plates, a putative dietary restriction treatment (Supplementary Figure 2); diverse manipulations of insulin/IGF-1 signaling (IIS), affecting the *daf-2* insulin/IGF-1 receptor and the *daf-16* FOXO transcription factor (Supplementary Figures 3-5), and manipulations of protein synthesis and mitochondrial function (Supplementary Figure 6) and the mTOR pathway (Supplementary Figure 7). As expected, interventions varied in the severity of their effects on senescent pathology. More notably, the severity of suppression differed between pathologies, with greater impacts often seen in the intestine and the pharynx, and weaker effects on uterine tumors; for details of effects of individual treatments, see Supplementary Information. The presence of such variation provides a useful basis for the application of ML to investigate pathology-lifespan relationships.

### Differential correlations between pathologies and metrics of declining health

That the age changes in anatomy documented here are deteriorative is largely self evident, hence their description as pathological^5, 29^. Other possible criteria for identifying them as pathological are that they contribute to age-related decline in health and to late-life mortality. We first probed the former possibility.

As *C. elegans* hermaphrodites grow older they exhibit various behavioral impairments affecting, among other things, locomotion, defecation and pharyngeal pumping. Age-changes in these three health-related parameters were measured, and the patterns of decline seen were comparable to those previously described (Figure 2a-c)^30, 31, 32^.

**Figure 2:**
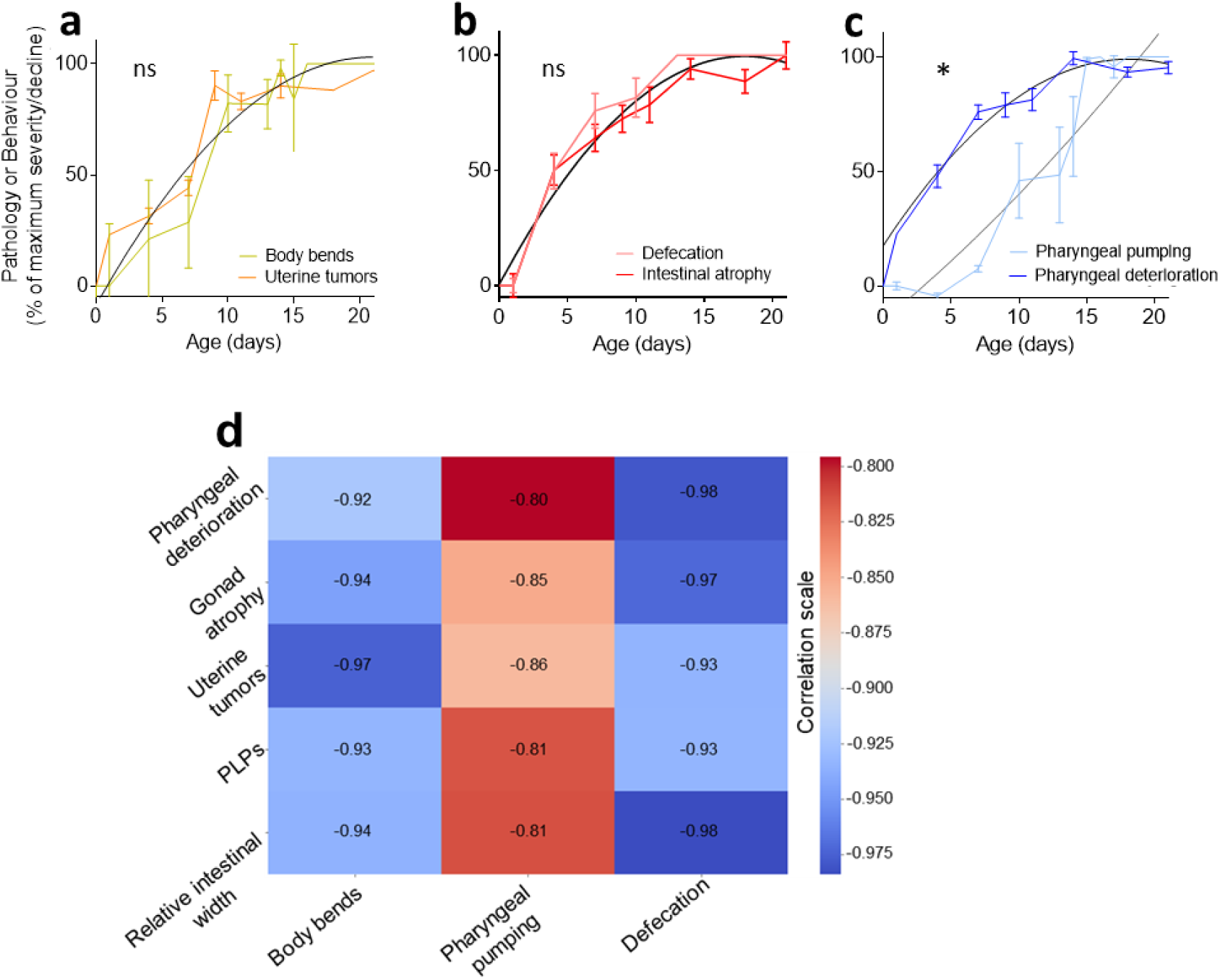
Variation in senescent pathology with health metrics. Health metric declines in early-mid adulthood. Selected comparison of fits (extra sum-of-squares F test). (a) Close correspondence of decline in movement (body bends) with uterine tumor progression, and (b) of decline in defecation with intestinal atrophy. (c) Loss in pharyngeal pumping occurs later than the major burst of pharyngeal pathology. 2 trials (*n* = 10). (d) Correlation matrix of the different pathologies measured and decline in health metric (analysis of raw data). Pearson method. **** *p* < 0.00001; *** *p* < 0.0001; ** *p* < 0.001; * *p* < 0.01.

Figure 2a-c displays comparisons of age changes in three pairs of health metrics and pathologies, while Figure 2d shows a correlation matrix between health and pathology metrics. Consider first the decline in locomotion. This proved to be most strongly correlated with uterine tumor development (Figure 2a, d). One possibility is that this reflects physical obstruction of body wall muscles by tumors, which frequently grow to fill much of the body cavity in the mid-body region.

If health is impaired by the hermaphrodite mid-life pathogenetic burst (Figure 1c), then its decline should coincide with the burst. This is in fact the case not only for locomotion, but also defecation, whose decline was particularly correlated with intestinal atrophy, and also pharyngeal deterioration and gonad atrophy (Figure 2a,b,d). By contrast, pharyngeal pumping declined some time later than the burst (including the appearance of pharyngeal pathology) (Figure 2c). This could imply that age changes in neuronal control of pumping rather than organ degeneration underlie the age-related decline in feeding rate. The coincidence of the mid-life anatomical changes and decline in movement and defecation provides further support for the designation of such changes as pathological.

### Negative correlation between senescent pathology development rate and lifespan

Next we investigated the relationship between senescent pathology and lifespan. To begin with we conducted an overall analysis of the correlation between lifespan and the average severity of all five pathologies. As a metric of pathology severity, pathology development rate was used. Rates were transformed into Z-scores (which describe a value’s relationship to the mean of a group of values) to allow pathologies to be compared to one another. This revealed, as one might expect, a negative correlation between mean pathology development rate and lifespan, both for *C. elegans* across all treatments, in individual wild-type worms followed throughout life, and in the other nematode species (R^2^=0.5; Supplementary Figure 8a). This is consistent with the view that senescent pathology contributes to late-life mortality.

### Use of machine learning to identify potential life-limiting pathologies

Results in this study underscore how different treatments that extend lifespan can suppress different senescent pathologies to differing extents. This raises the possibility that different combinations of pathologies may affect lifespan to different degrees, and in an independent or dependent manner (i.e. one pathology may influence the effect on lifespan of another pathology). A further possibility is that the relationship between pathology severity and lifespan differs between pathologies. To investigate the complex relationship between pathology severity and lifespan, an ML-based approach was employed. To obtain a dataset to interrogate that was of a sufficiently large size for ML analysis, we combined data generated in this study with various data from previous studies, from our lab and others. This included data on effects of a variety of mutations, RNAi treatments and culture condition variations (e.g. in temperature, and monoxenic vs axenic culture) scored at the population level (i.e. population average metrics), along with data from wild-type animals scored at both the individual and population levels. For details of all data sources and conditions see Supplementary Table 2.

First, we trained and evaluated different standard models (linear regression, ElasticNet regression, Support Vector Machine [SVM], random forest [RF], multilayer perceptron [MLP]) to roughly identify the best predictor of lifespan based on raw pathology scores. The random forest (RF) model outperformed the others (R^2^=0.57; mean average error [mae] of predicted life of 4.0 days)(Supplementary Figure 8b).

Next, to account for the fact that the pathologies are progressing with time and that pathology values at different time points are not independent of one another, differences in scores through time were fed into the different models (i.e. pathology scores at each time point had the previous time point scores subtracted). Again, the RF model outperformed the others, yielding an R^2^ of 0.79 and mae of 2.77 days; and an average R^2^ of 0.7 and mae of 2.6 days on unseen evaluation sets (20 runs) (Figure 3a). In other words, >70% of the variation in nematode lifespan between different conditions and treatments can be predicted using the five senescent pathologies characterized here.

**Figure 3.**
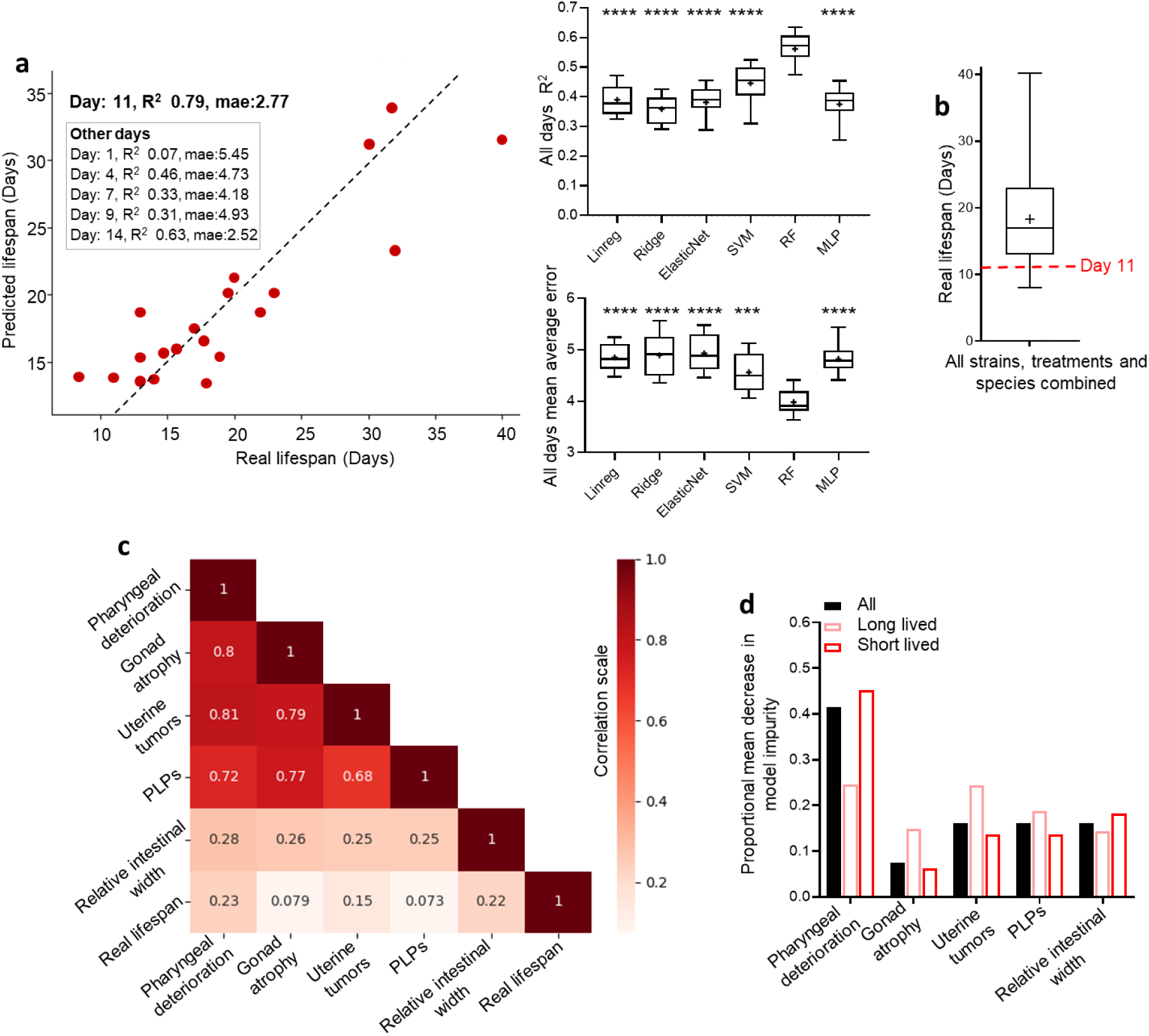
Machine learning can predict lifespan from senescent pathology. Analysis here used a random forest (RF) ML algorithm and includes all animals (single wild-type worms, treatments and species). (a) Left: scatter dot plot showing real observed lifespan against lifespan predicted by the RF ML model. The data was randomly split with 80% data used to build the model, and the remaining 20% used to test the model and generate the data displayed. Right: R^2^ and mean average error (days) with different AI models (linear regression, ElasticNet regression, Support Vector Machine (SVM), random Forest (RF), multilayer perceptron (MLP) independent of age at which the pathology was measured (i.e. pathology progression through time). For models using only data up to day 11 of adulthood see Supplementary Figure 8b. For coefficients of the linear regression model see Supplementary Figure 8c. pval comparing RF model to other models. (b) All wild-type single animal lifespans, and average population lifespans of various treatments and species used to build the model plotted as a box plot. The average lifespan of all single wild-type worms, treatments and species is 18.3 days. Note that some treatments extend lifespan while others shorten lifespan. Day 11 (shown in red), the time point up to which pathology progression was scored and used to predict lifespan in (a). (c) Correlation matrix of the different pathologies measured (raw data) and the real observed lifespan. Pearson method. (d) Feature importance: RF model Mean Decrease in Impurity; long lived refers to worms having survived for more than 30 days. s.e.m. displayed following 20 model runs; *t*-test: long lived vs short lived animals. * *p* < 0.01, ** *p* < 0.001, *** *p* < 0.0001, \*\*\*\**p* < 0.00001. For specificity and sensitivity analysis based on predicting long-lived vs short-lived animals see Supplementary Figure 8d. For feature importance of individual pathologies relative to mutant animals (and animals in non-standard culture conditions) vs wild type see Supplementary Figure 8e. For raw pathology and lifespan data used see Supplementary Table 2.

Also, in the RF model pathology progression up to day 11 performed better than progression up to day 14; this likely reflects the fact that the severity of most pathologies plateaus after day 11 (Figure 1c)^5^. For reference, the mean lifespan of all single wild-type animals and treated animal populations used in this study is 18.3 days (lower quartile: 13 days; Figure 3b).

In terms of model feature importance, the pharynx and intestinal pathology scores correlate the most strongly with observed lifespan (Figure 3c). Furthermore, the greatest determinant of model prediction (based on Gini index or Mean Decrease in Impurity [MDI]) is pharyngeal pathology, by a wide margin (Figure 3c). This is consistent with previous findings pointing to links between lifespan and both pharyngeal and intestinal status^5, 18, 33, 34^.

Next we looked at differences between long- and short-lived single wild-type worms, as well as long- and short-lived treatment populations and species used in the model. Here long lived was defined as >18 days and short lived <18 days. This value was selected as it is the approximate mean *C. elegans* lifespan at 20°C as well as the mean of all single wild-type worms, treatments (culture under standard vs non-standard conditions, e.g. axenic media) and species used in this study (Figure 3b).

This analysis reveals that pharyngeal pathology is more predictive of lifespan in short-lived animals than in long-lived ones (Figure 3d). One possibility is that this reflects increased early deaths linked to pharyngeal infection^33^. By contrast, intestinal atrophy and PLPs are similarly predictive of life in short- and long-lived animals. Notably, uterine tumors became more predictive of lifespan with treatments that extend life. This suggests that in animals subjected to life-extending treatments, the contribution of uterine tumors to late-life mortality increases.

### Early burst of pathogenesis is absent from *C. elegans* males

Finally, we describe one notable finding from our initial survey of senescent pathologies in different contexts. *C. elegans* can reproduce either as self-fertilizing hermaphrodites or by mating with males. Sex differences in several individual pathologies have been previously noted, as follows. Aging virgin males do not accumulate yolk pools in the body cavity^5^ consistent with the absence of vitellogenin (yolk protein) synthesis in males^35^. Moreover, virgin males do not exhibit distal gonad atrophy^36^ or marked intestinal atrophy^5^.

To give a fuller picture of sex differences in senescent pathology, we directly compared all five pathologies in the two sexes (unmated) (Figure 4a-e). This confirmed the absence of yolk pools, intestinal atrophy and gonad atrophy in virgin males. Moreover, we detected no form of germline tumor equivalent to the teratoma-like uterine tumors seen in senescent hermaphrodites^16, 37^. By contrast, similar levels of pharyngeal pathology were seen in the two sexes (Figure 4a-e). Thus, major pathologies develop rapidly in organs linked directly to reproduction (i.e. the gonad and intestine) in hermaphrodites but not males. This could imply that trade-offs linked to reproduction lead to pathology in hermaphrodites but not males. These results demonstrate major sexual dimorphism in senescent pathophysiology in *C. elegans* and therefore, potentially, in the causes of late-life mortality. They also hint at a greater role of pharyngeal pathology in late-life mortality in males than in hermaphrodites.

**Figure 4:**
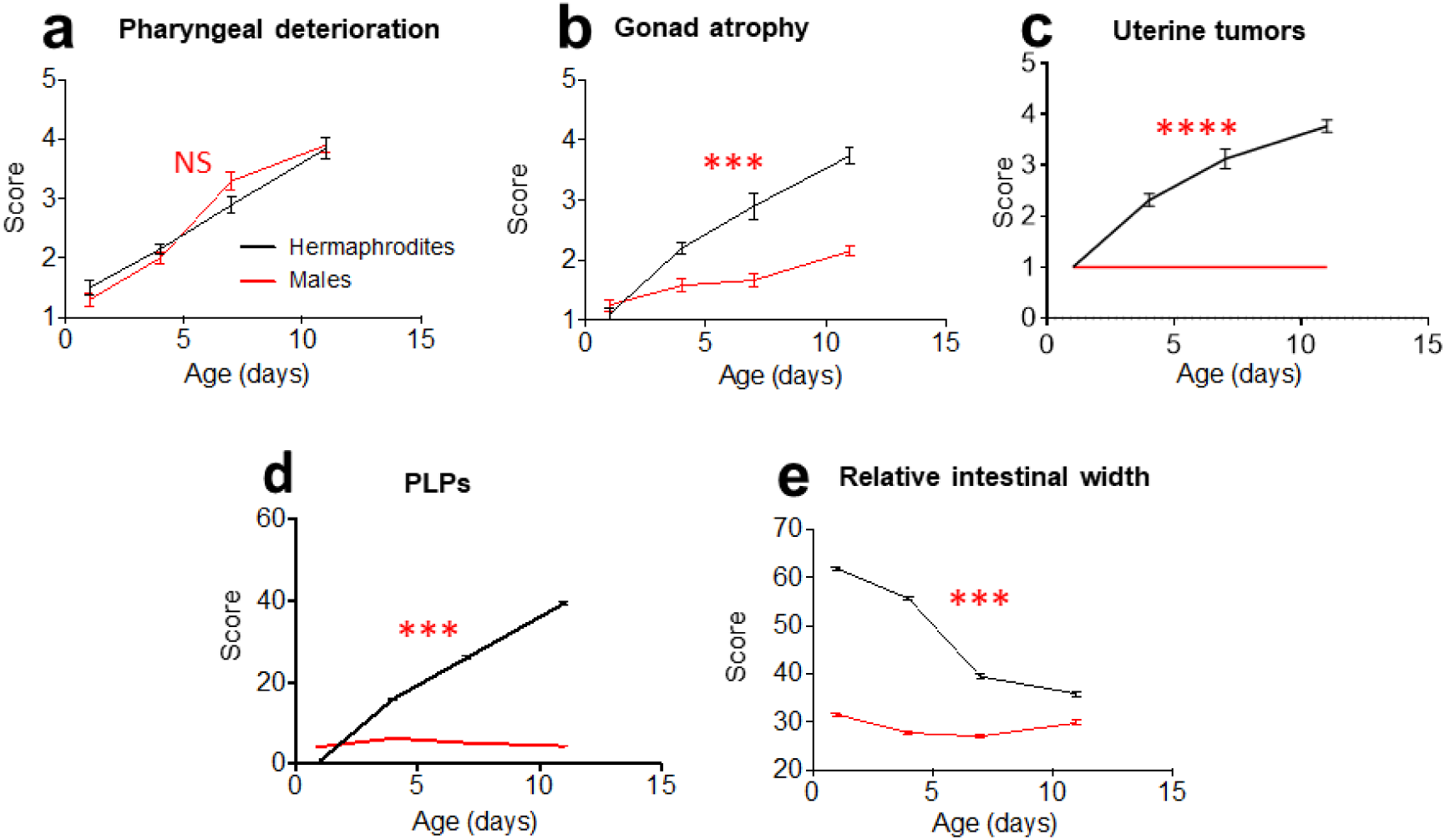
Sex difference in senescent pathology. Most of the age-related pathologies seen in hermaphrodites are largely absent from males (two-way ANOVA, Bonferroni correction), and stars show statistically significant differences in pathology progression (ANCOVA, Tukey correction). (a) Pharyngeal deterioration is similar in both sexes. (b) Gonad atrophy is absent from males. (c) Germline tumors (teratoma-like uterine tumors in hermaphrodites) and (d) PLPs (yolk pools) are absent from males. (e) Gut atrophy is largely absent from males. (f-h) Health metric declines in early-mid adulthood. Selected comparison of fits (extra sum-of-squares F test). **** *p* < 0.00001; *** *p* < 0.0001; ** *p* < 0.001; * *p* < 0.01.

## Discussion

In this study we have described how effects of interventions that alter lifespan have differential effects on senescent pathology. Capitalizing on this, we have investigated the relationship between pathology and lifespan using a ML approach. This suggested pharyngeal and intestinal pathology as greater causes of late-life mortality. In life-shortening contexts the importance of pharyngeal pathology increased, as did that of uterine tumors in life-extending contexts.

### Use of ML analysis: insights and limitations

The use of several ML models to predict lifespan from pathology were tested and one, the random forest (RF) model, consistently outperformed the others. Our analysis shows that non-linear ML models consistently outperformed the linear ones when predicting lifespan. In particular, and interestingly, RF regression models yielded the best performance. A possible reason for this is the contingent nature of the relationship between pathology and survival. Rather than linear or additive relationships, it may be that given pathologies only limit lifespan at specific severities and in specific combinations with one another. We suggest that it may be for this reason that creation of decision trees by the RF model has the greatest predictive power. (Decision trees are where the model learns how to best split the dataset into smaller and smaller informative subsets to predict the target value).

The models are able to predict 70% of the variation in lifespan, but what about the remaining 30%? Here it should be noted that our dataset included data compiled from other studies where minor differences in culture conditions could create noise. For example, the mean lifespan of *C. elegans* under standard conditions at 20°C typically ranges from 17 to 21 days. Moreover, pathology is typically scored only every 3-5 days. Furthermore, this study was limited to five more easily assayed senescent pathologies, and other, less visible pathologies are very likely to contribute to mortality. Not included here are, for example, muscle atrophy, cuticular hypertrophy^6^ and senescent changes in the nervous system such as neurite outgrowths, synapse deterioration^38, 39^ and PVD neuron degeneration^20^.

### Is lifespan a function of overall aging or specific life-limiting pathologies?

The correlation between intestinal pathology and lifespan supports a central role the intestine in nematode aging. This could be because the gut regulates aging in the whole organism, by activating signalling pathways in other tissues, or because gut pathology itself is life limiting. Notably, DAF-16 activity in the gut partially restores *daf-2* mutant longevity in *daf-16; daf-2* mutants. There is evidence that this involves paracrine action via both DAF-16-dependent and -independent responses, involving alteration of gene expression^40, 41, 42^ and of various processes in other tissues^43, 44, 45^, reviewed by^46^. Thus, the intestine may regulate aging in the whole organism via paracrine (and also autocrine) signaling.

An further possibility is that intestinal pathology also limits lifespan. Limitation of life by organ-specific pathology is a plausible explanation for the correlation of pharyngeal deterioration with mortality, given that suppression of early death from pharyngeal *E. coli* infection increases lifespan^33, 34, 47^. If intestine-limited rescue of *daf-16* in *daf-16; daf-2* double mutants extends lifespan mainly by regulating whole-animal aging, one would expect to see amelioration of multiple pathologies. By contrast, if it acts by preventing life-limiting pathology in the gut, we would expect little effect on other pathologies. In fact, the latter was the case (Supplementary Figure 5), suggesting that intestinal pathology contributes to late-life mortality. Thus, life-extension from intestinal rescue of *daf-16* (Ref 40) and of other factors^5, 48^ could reflect, at least in part, rescue of life-limiting intestinal pathology.

### A context-dependent role of uterine tumors in late-life mortality?

Could uterine tumors contribute to late-life mortality? While metastatic cancer is a major cause of late-life mortality in mammals, the mammalian teratomas that *C. elegans* uterine tumors resemble are usually benign^16^. Consistent with this, interventions that selectively block tumor growth do not increase lifespan in otherwise wild-type populations^17, 49^. Moreover, we observed that mutation of *daf-2*, that greatly extend lifespan, suppresses several pathologies, but not uterine tumors (Supplementary Figure 3), arguing against a life-limiting role. However, the greater correlation between uterine tumor suppression and longevity in long-lived contexts revealed by ML analysis suggests that this pathology could become life-limiting in such circumstances. One possibility is that amelioration of pathologies that limit life leads to their replacement by new causes of late-life mortality. This warrants further investigation.

### Sexual dimorphism in pathology development and aging

The aging process exhibits sexual dimorphism in a number of ways, ranging from sex differences in evolutionary determinants of aging^50^, in effects of interventions that increase lifespan^51^, and in patterns of aging and late-life disease in men and women^52, 53^. This is true of *C. elegans* too: for example, mutation of the *daf-12* dafachronic acid receptor alters lifespan in hermaphrodites^54^ but not males^55^, while some mutations that impair neuronal synapse function increase lifespan in males but not hermaphrodites^56^.

The present study reveals profound sex differences in senescent pathology: the severe age-related pathologies seen in hermaphrodites are largely absent in males, with the exception of pharyngeal deterioration (Figure 4a-e). This demonstrates fundamental differences in the aging process between *C. elegans* hermaphrodites and males. Coincident with their reduced pathology, the lifespan of wild-type males is typically about 20% longer than that of hermaphrodites, at least when life-shortening male-male interactions are prevented^55, 56, 57, 58^. Possibly this reflects the reduced levels of senescent pathogenesis in males, apart from that affecting the pharynx. This could imply that senescence of the pharynx plays a greater role in late-life mortality in males.

Recent evidence suggests that the severe pathologies in hermaphrodites could reflect the occurrence of semelparous reproductive death^18, 23, 25^. The absence of this pathology in males implies that reproductive death in *C. elegans* is hermaphrodite specific. Consistent with this, prevention of reproductive death in semelparous species can result in large increases in lifespan^18^, and in *C. elegans* germline removal (which blocks senescent pathology in hermaphrodites) increases lifespan in hermaphrodites^59, 60^ but not males^61^.

### A two-stage model of programmatic aging in *C. elegans* hermaphrodites

The ultimate aim of the work in this study is to understand the causes of senescence (aging) and the etiologies of senescent pathology across the animal kingdom (including humans). For many decades, aging has been viewed by biologists as a passive process of breakdown, wearing out and damage accumulation^62, 63, 64, 65^. By contrast, the senescent pathologies described in this study appear more the result of genetically determined, wild-type processes that actively generate pathology in a programmatic fashion.

These processes could on the one hand reflect futile run-on of processes that promote reproductive fitness in early adulthood, as in the run-on of embryogenetic programs in unfertilized oocytes that generates the abnormal, hypertrophic cells of uterine tumors^16, 37^. On the other hand, they could reflect costly programs which promote fitness, as appears to be the case for intestinal atrophy coupled to production of yolk that is vented through the vulva to support larval growth in a milk-like fashion^5, 23, 24^.

Thus, the self-destructive mechanisms of senescent pathophysiology are healthy, vigorous processes, consistent with the hypothesis that hyperfunction can generate pathology through programmatic mechanisms^4, 29, 66, 67, 68^. It is notable that a phase of rapid and extensive anatomical change (day 4-12) is followed by a period of relative anatomical stasis^5^. This suggests the following hypothesis about aging in *C. elegans* hermaphrodites: that it occurs in two stages. First, a stage of hyperfunction (days 4-14), where the organism vigorously destroys its internal organs (particularly the alimentary canal and reproductive system). Second, a resulting second, terminal state of profound physiological impairment in which the organism is, effectively, mortally ill. The duration of subsequent survival (days 15-21) represents the time that it takes for the worms to die as a consequence of prior pathogenetic changes (Figure 5).

**Figure 5.**
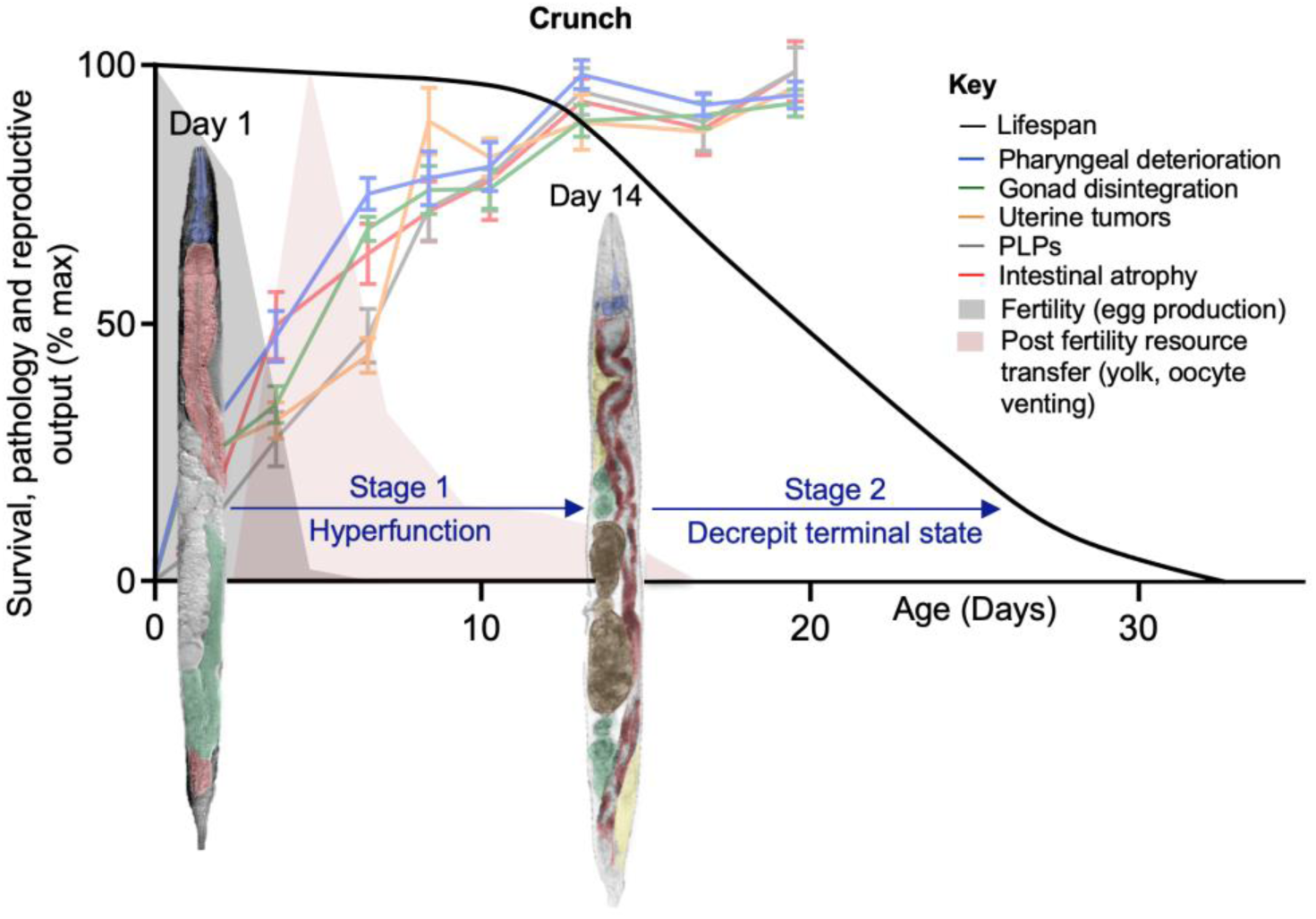
A two-stage model of programmatic aging in *C. elegans* hermaphrodites. Here the process of senescence begins with a first phase in which active, programmatic processes drive rapid and extensive anatomical change (hyperfunction; days 4-14). This period ends as these processes cause a general breakdown of physiology (the crunch). Thereafter, nematodes linger on in a decrepit state of terminal illness from which they gradually die. Production of progeny largely ends by day 5, but for some days after that resource transfer to progeny can occur by means of yolk venting^24^.

## Methods

### Culture methods and strains

*C. elegans* were maintained at 20°C on Nematode Growth Medium (NGM) plates seeded with *Escherichia coli* OP50, unless otherwise stated. 5-fluoro-2-deoxyuridine (FUDR), sometimes used to block progeny production, was not used in this study. *C. elegans* strains used in this study included: N2M male stock *fln-2(ot611) X (*Ref. 47), CF1724 *daf-16(mu86) I; daf-2(e1370) III; muIs105 [daf-16p::GFP::daf-16 + rol-6(su1006)]*, CF2005 *daf-16(mu86) I; daf-2(e1370) III; muIs120 [ges-1p::GFP::daf-16 + rol-6(su1006)],* CF2093 *daf-16(mu86) I; daf-2(e1370) III; muIs131 [unc-119p::GFP::daf-16 + rol-6(su1006)]*, CF2102 *daf-16(mu86) I; daf-2(e1370) III; muIs126 [myo-3p::GFP::daf-16 + rol-6(su1006)]*, DR1563 *daf-2(e1370) III*, DR1572 *daf-2(e1368) III,* GA82 *daf-2(e1370) III,* GA114 *daf-16(mgDf50) I*; *daf-2(e1370) III,* GR1307 *daf-16(mgDf50) I,* MQ887 *isp-1(qm15) IV,* MQ1333 *nuo-6(qm200) I* and RB1206 *rsks-1(ok1255) III*.

Pathology and survival data used in ML analysis was drawn both from this study, and a number of earlier studies. For the latter, strains and sources are listed in Supplementary Table 2.

### Preparation of aging cohorts for pathology measurements

For each condition, approximately 200 animals were aged on 4 different plates, starting from the pre-adult L4 stage. Animals were transferred to fresh plates daily during reproduction (to avoid overgrowth with offspring) and at approximately weekly intervals thereafter. In trials using RNA interference (RNAi), HT115 RNAi-producing (clones *ife-2*, *iftb-1*, *hsf-1*) and control bacteria (plasmid L4440), were cultivated as previously described^69^, washed and seeded onto NGM plates. RNAi treatment was started at the L4 stage. For trials involving males, animals were maintained at low population density (∼5 per plate) to reduce detrimental effects of male-male interactions^56^ and transferred to fresh plates at the same time as hermaphrodites.

### Pathology measurements

Animals were imaged on days 1, 4, 7, 11, 14 and 18. At each time point, 10-15 animals were mounted onto 2% agar pads and anesthetized with 0.2% levamisole. DIC images were acquired with an Orca-R2 digital camera (Hamamatsu) and either a Leica DMRXA2 microscope or a Zeiss Axioskop 2 plus microscope, driven by Volocity 6.3 software (Improvision, Perkin-Elmer). Images of pathology were analysed semi-quantitatively^5, 7, 17^ (Table 1, Figure 1).

For pharynx, gonad and tumor pathologies, images were randomized, examined by trained scorers, assigned severity scores of 1-5, and mean values calculated. Here 1 = youthful, healthy appearance; 2 = subtle signs of deterioration; 3 = clearly discernible, mild pathology; 4 = well-developed pathology; and 5 = tissue so deteriorated as to be barely recognizable (e.g. gonad completely disintegrated), or reaching a maximal level (e.g. large uterine tumor filling the entire body width). Intestinal atrophy was quantified by measuring the intestinal width at a point posterior to the uterine tumors, subtracting the width of the intestinal lumen and dividing by the body width. Yolk accumulation was measured by dividing the area of pseudocoelomic yolk pools by the area of the body visible in the field of view as captured at 630x magnification.

Single worm pathology measurements were also performed throughout life. Animals were mounted onto 2% agar pads on a microscope slide and the slides placed on ice for 5 min. Animals were then immediately imaged and recovered. Animals were imaged on days 1, 4, 7, 11, 14, and 18. Lifespan was also measured in each animal.

### Axenic culture

For experiments using axenic medium plates, with no *E. coli*, plates were prepared as previously described^70^. Briefly, gravid hermaphrodites were bleached and eggs plated on NGM plates bearing UV-irradiated *E. coli*, to allow normal growth to the L4 stage. L4 larvae were then transferred to and maintained upon axenic medium plates.

### Statistical analysis

Pathology development was analysed using two-way ANOVA to compare different time points. Differences in pathology progression were compared using an ANCOVA, with Z-scores for standardization of different pathologies, as described^18^. Bonferroni’s or Tukey’s tests were used to correct for multiple comparisons.

### Machine learning model generation

Machine Learning (ML) regression models were trained to predict the lifespan of nematodes (individual wild-type animals under standard conditions, or populations with altered genotypes/culture conditions) based on pathology development features. For individuals, wild-type worms were followed throughout life and day of death was scored as well as their pathology levels on days 1, 4, 7, 11, and 14. For populations, pathology was scored in samples of 10 individuals at each time point, and mean lifespan of the population was used.

More precisely, ML models were trained to learn a mapping *f*: *X* → *y*. Here *X* = [*x*_1_ … *x*_*N*_*F*__] representing the observed pathology development features (number of features: *N*_*F*_ = 5): pharyngeal deterioration, gonad atrophy, uterine tumors, PLPs and intestinal width, observed at each time point. *y* represents the lifespan of a particular worm individual or population. In total, 434 observations were used to train and evaluate the ML models. A small percentage (<10%) of pathology development values were missing due to measurement and recording errors (e.g. where captured images were too unclear to score pathology). These missing values were replaced with the mean values of the pathology feature observed at a specific day. For example, the missing pharyngeal deterioration value of a worm observed at day 7 was set to the average of all pharyngeal deteriorations observed at day 7. The target value *y* for the ML model predictions was the observed lifespan of the worm minus the day at which the pathology features were observed.

Three linear ML models (Linear Regression, Ridge Regression and ElasticNet regression) and three non-linear models (Support Vector Machine [‘rbf’ kernel], Random Forest [500 estimators], and a Multilayer Perceptron with one hidden layer ([5 hidden nodes, Tanh activation function]) were evaluated. The models were implemented in Python using the scikit-learn toolbox^71^. Each model was trained and evaluated with 20 random train-test (80:20) splits of the initial data.

## Supplementary information

Supplementary data are available at. Files include: Supplementary Information; Supplementary File 1 of raw data used to build the model.

## Acknowledgements

We thank Jennifer Tullet (University of Kent) and members of the Gems lab for useful discussion and/or comments on the manuscript. Some strains were provided by the Caenorhabditis Genetics Center, which is funded by NIH Office of Research Infrastructure Programs (P40 OD010440).

## Author contributions

M.E., D.G. and C.C.K. contributed to wet lab project design. C.C.K developed theory behind mathematical and AI/ML analysis and performed results interpretation. C.A., M.E., A.F.G., C.C.K., H.W. and A.Z. performed wet lab data collection. M.C., and P.M.S. performed AI/ML data tabulation and analysis. D.G. initially drafted the manuscript, and M.E., D.G. and C.C.K. then edited the manuscript.

## Supplementary Information

### Contents Summary

#### Other supplementary files

**Supplementary File 1.** All raw pathology and lifespan data.

**Supplementary File 2.** AI code used to generate model.

### Supplementary results

#### Effects of axenic culture on senescent pathology

We examined how interventions that alter lifespan affect the rate of development of senescent pathology. Dietary restriction (DR) increases lifespan and delays senescent pathology in a number of organisms^1^. Several putative *C. elegans* DR regimens have been described^2^. One such is culture on nutrient-rich, semi-defined axenic medium (i.e. lacking a microbial food source), which causes retarded development and large increases in lifespan and therefore resembles a DR effect^3,4^. We employed a solid medium protocol where larvae are raised on UV-killed *E. coli* OP50 to support normal growth and then transferred onto axenic medium plates (no *E. coli*) at adulthood^5^. This resulted in almost complete suppression of pharyngeal deterioration, PLPs and intestinal atrophy, and partial suppression of uterine tumors (Supplementary Figure 2a-d). Gonad atrophy was not measured as axenic culture inhibited gonadal development.

#### Selective effects on pathology of different *daf-2* alleles

Reduction of insulin/IGF-1 signaling (IIS), as in *daf-2* insulin/IGF-1 receptor mutants, greatly increases *C. elegans* lifespan, and this effect is dependent upon the DAF-16 FOXO transcription factor^6^. Mutation of *daf-2* suppresses development of diverse pathologies, including gonadal and pharyngeal deterioration^7^, late-life bacterial infection in the pharynx ^8,9^, intestinal atrophy and yolk accumulation^10,11^. Inhibiting bacterial infection or yolk synthesis is sufficient to extend lifespan^12-14^, suggesting that reduced IIS increases lifespan by blocking development of pathology, at least partly.

We previously compared effects on intestinal atrophy and yolk pool accumulation of two *daf-2* mutations, a less pleiotropic (class 1) allele *daf-2(e1368)*, and a more pleiotropic (class 2) allele *daf-2(e1370)*^15^. This showed that *e1368* modestly delayed both pathologies, but *e1370* strongly suppressed them^11^. Here we extend this analysis to the remaining three pathologies. Notably, the five pathologies were not regulated by IIS as a group, in tandem with one another. For example, in *e1370* mutants, development of pharyngeal and gonadal pathology was delayed but eventually reached maximal wild-type levels, whereas for intestinal atrophy and yolk pools they did not. Moreover, little reduction in uterine tumor growth was detected in either mutant (Supplementary Figure 3a-e). The lack of effect was also somewhat unexpected, given the previous observation that *daf-2* also inhibits formation of chromatin masses within uterine tumors^16^; it suggests that *daf-2* reduces chromatin mass formation without reducing tumor size.

All suppression of pathology by *daf-2(e1370)* was fully dependent upon *daf-16* FOXO, as is the case for *daf-2* longevity^17^, consistent with the possibility that IIS shortens lifespan by causing senescent pathology (Supplementary Figure 3f-j).

These findings imply that although wild-type IIS promotes all five senescent pathologies, it does not do so equally. Rather, IIS strongly promotes intestinal atrophy and yolk accumulation, modestly promotes gonadal atrophy and pharyngeal deterioration, and has little effect on uterine tumor formation. Also, different alleles differ in terms of severity of suppression of pathology as well as mortality: *e1370* has stronger effects than *e1368* on both pathology (including pharyngeal infection, gut atrophy and PLPs) and lifespan^8,9,11^.

Next we investigated the role of the *daf-16* FOXO transcription factor gene in development of senescent pathology. Mutation of *daf-16* slightly shortens lifespan^18^ and fully suppresses *daf-2* longevity^17^. The null mutation *daf-16(mgDf50)* reduced pharyngeal deterioration slightly and did not significantly affect gonad atrophy or tumor formation, but did aggravate PLPs and intestinal atrophy (Supplementary Figure 4a-e). This is in agreement with an earlier study showing little effect of *daf-16* on the pharynx and germline^7^; and our observation that *daf-16(0)* accelerates PLPs and intestinal aging^11^.

We also examined the effects of the *hsf-1* heat shock factor, another transcription factor that mediates the effect of *daf-2* on lifespan^19^. It was previously shown that *hsf-1* RNAi substantially shortens lifespan and accelerates pharyngeal pathology^7^. Although we could not confirm the effects of *hsf-1* RNAi on pharyngeal pathology, we did observe accelerated gonad atrophy and intestinal atrophy (Supplementary Figure 4f-j). These results could imply that loss of *daf-16* or *hsf-1* shortens lifespan by promoting pathology development, and support the earlier deduction that *hsf-1* RNAi causes a progeroid condition^7^.

Next we examined whether effects of *daf-2(e1370)* on pathology are *daf-16* dependent, and found that they are, fully (Figure 4f-j). This raises the question of where, within *daf-2(e1370)* mutants, *daf-16(+)* acts to prevent pathology development. With respect to lifespan, effects of *daf-16* are exerted most strongly in the intestine, as shown using transgenic strains where *daf-16* is expressed from tissue-specific promoters in a *daf-16; daf-2* background^20^. We used the same strains to ask: is this also true of the action of *daf-16* against senescent pathologies?

Expressing *daf-16* with its own promoter restored the capacity of *daf-2(e1370)* to suppress pathology in all four cases, confirming the anti-pathogenic role of *daf-16* (Supplementary Figure 5a-f). In most cases, *daf-16* expression in muscle or in the nervous system had little effect on any pathologies (Supplementary Figure 5a-f). By contrast, intestinal rescue of *daf-16* partially restored *daf-2* suppression of intestinal atrophy and yolk steatosis, suggesting an organ autonomous effect of *daf-16*. Intestine-limited rescue of *daf-16* did not suppress gonadal or pharyngeal pathology, suggesting that *daf-16* acts in a tissue- or organ-autonomous manner to suppress pathology.

That *daf-16* rescue in the intestine suppressed both intestinal atrophy and PLP formation (Supplementary Figure 5d,e) is consistent with tissue autonomous suppression of conversion of intestinal biomass into yolk by *daf-16*^11^. By contrast, intestinal atrophy is also modestly suppressed by neuronal rescue of *daf-16* (Supplementary Figure 5d,e), implying tissue non-autonomous action of *daf-16*. Notably, neuronal (as well as intestinal) knockdown of DAF-2 auxin-induced DAF-2 degradation increases lifespan^21-23^ while neuronal rescue of *daf-2* shortens it^24^; possibly intestinal *daf-16* effects on biomass conversion play a role in this.

#### Effects on senescent pathology of other interventions that extend lifespan

Next we investigated effects on senescent pathology of life-extending genetic interventions that inhibit protein synthesis, the mTOR pathway and mitochondrial function. Knockdown of genes involved in protein translational machinery can increase lifespan, including the initiation factor 4F (eIF4F), *ife-2*, the translation initiators eIF2β (*iftb-1*) and eIF4G (*ifg-1*), and the small ribosomal subunit *rps-15*^25^. We found that *ife-2* and *iftb-1* RNAi had little effect on pharyngeal pathology, gonad atrophy or uterine tumors. By contrast, both interventions suppressed intestinal atrophy, particularly *ife-2* RNAi (Supplementary Figure 6a-e). Possibly this reflects reduction of vitellogenin synthesis, which is coupled to intestinal atrophy ^11,14^. *iftb-1* RNAi also markedly increased PLP levels (Supplementary Figure 6d) which was somewhat unexpected; however, inhibition of *iftb-1* can reduce brood size by 50%^26^, which is predicted to cause yolk retention due to reduced efflux via egg laying^14^.

Protein synthesis is promoted by the target of rapamycin (TOR) pathway, acting via ribosomal protein S6 kinase (S6K), encoded by *rsks-1* in *C. elegans*. Lifespan is increased by loss of function of several genes in the TOR pathway, including *rsks-1(ok1255)*^27^, which also reduces protein synthesis rate^28^. *rsks-1(ok1255)* slightly alleviated pathology development, with significant effects on pharyngeal decline and tumor formation. Intestinal atrophy appeared delayed but the effect was not statistically significant (Supplementary Figure 7). The modest effects seen are consistent with the small magnitude of effects of *rsks-1* on lifespan, e.g. a 6% increase in mean lifespan in one study (20°C, *E. coli* HT115)^28^.

Finally, we tested mutations that perturb mitochondrial function and also delay development and increase lifespan, the Mit phenotype^29^, specifically point mutations in *isp*□*1*, encoding a catalytic subunit of mitochondrial complex III^30^ and *nuo-6*, a subunit of mitochondrial complex I^31^. *isp-1(qm150)* and *nuo-6(qm200)* both inhibited pathology development of most but not all of the five pathologies (Supplementary Figure 6f-j), indicating a major role of mitochondrial function in senescent pathogenesis.

One possibility is that mitochondrial metabolism supports processes that contribute to pathogenesis, such as feeding and reproduction. E.g., pharyngeal pumping contributes to pharyngeal pathology^32^, production of oocytes past the reproductive period results in uterine tumors^33^, and yolk synthesis results in PLP accumulation and intestinal atrophy^11^. Decreasing mitochondrial function reduces both pharyngeal pumping and reproduction^31,34,35^.

## Supplementary figures

**Supplementary Figure 1.**
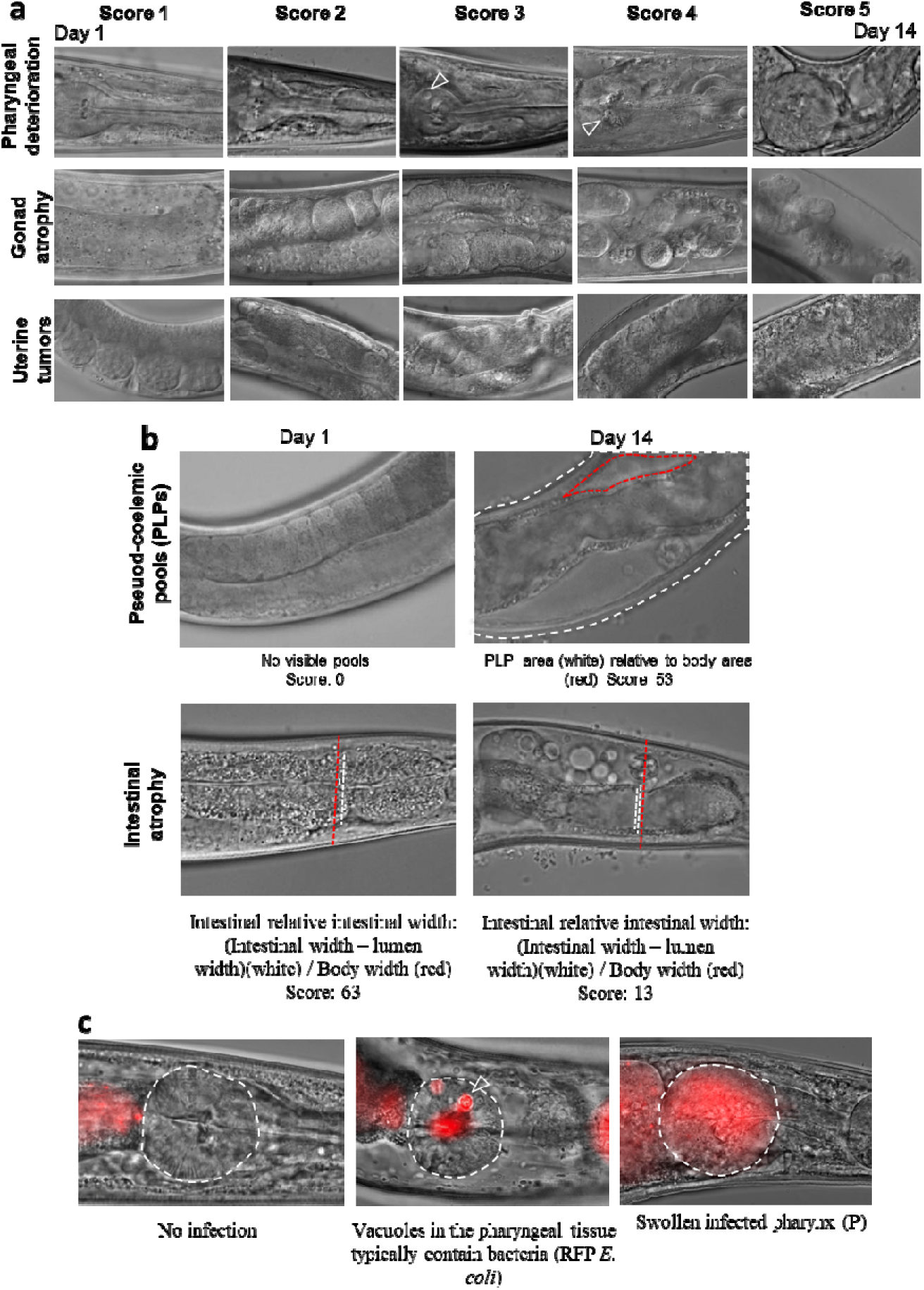
Quantification of age-related pathologies in aging *C. elegans.* (a) Images showing age-related pathologies in *C. elegans* hermaphrodites: Pharyngeal deterioration, gonad atrophy, uterine tumors, PLPs, and intestinal atrophy. Images were randomized, and given scores of 1-5. Here 1 = a youthful, healthy appearance; 2 = showing subtle signs of deterioration; 3 = clearly discernible, low level pathology; 4 = well developed pathology; and 5 = tissue so deteriorated as to be barely recognizable (e.g. gonad completely disintegrated), or reaching a maximal level (e.g. large tumor filling the entire body diameter). (b) Pseudocoelomic lipoprotein pool (PLP) formation (yolk accumulation) and intestinal atrophy. PLPs were measured by dividing the total area of yolk pools with the area of the body visible in the field of view. Intestinal atrophy was measured by calculating the width of intestinal tissue relative to body width. (c) Bacterial infection in the pharynx shown via RFP::*E. coli.* Arrow: vacuole site of infection. Terminal bulb of the pharynx is circled.

**Supplementary Figure 2:**
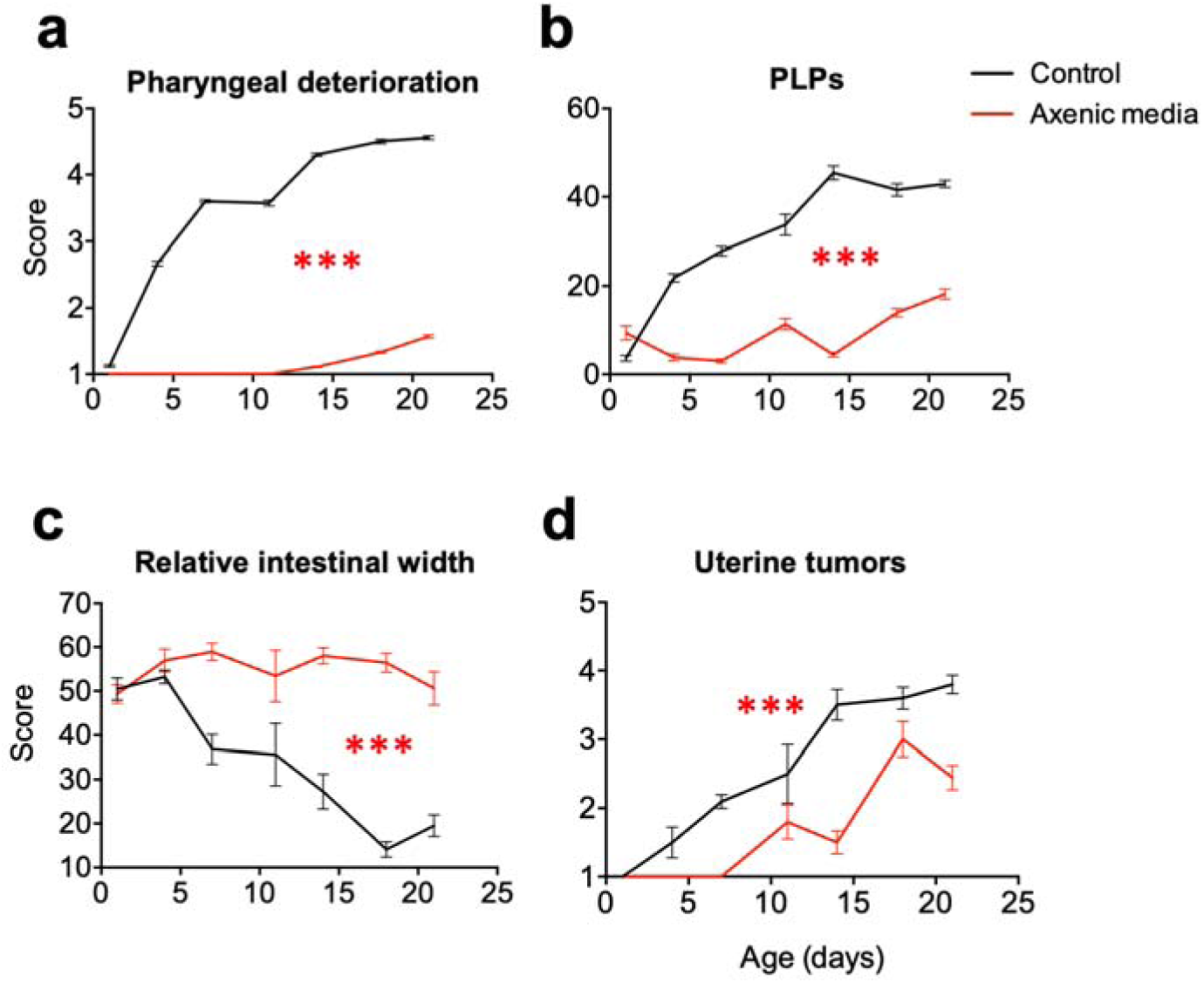
Effects on senescent pathologies of axenic culture (putative DR regimen). (a-d) Culture on solid axenic medium suppresses all age-related pathologies. (a) Pharyngeal deterioration, (b) PLPs, (c) gut atrophy, (d) uterine tumors. (a-d) 2 trials (*n*=10). Stars show differences in pathology progression from day 1 to 14; ANCOVA (Tukey correction).

**Supplementary Figure 3.**
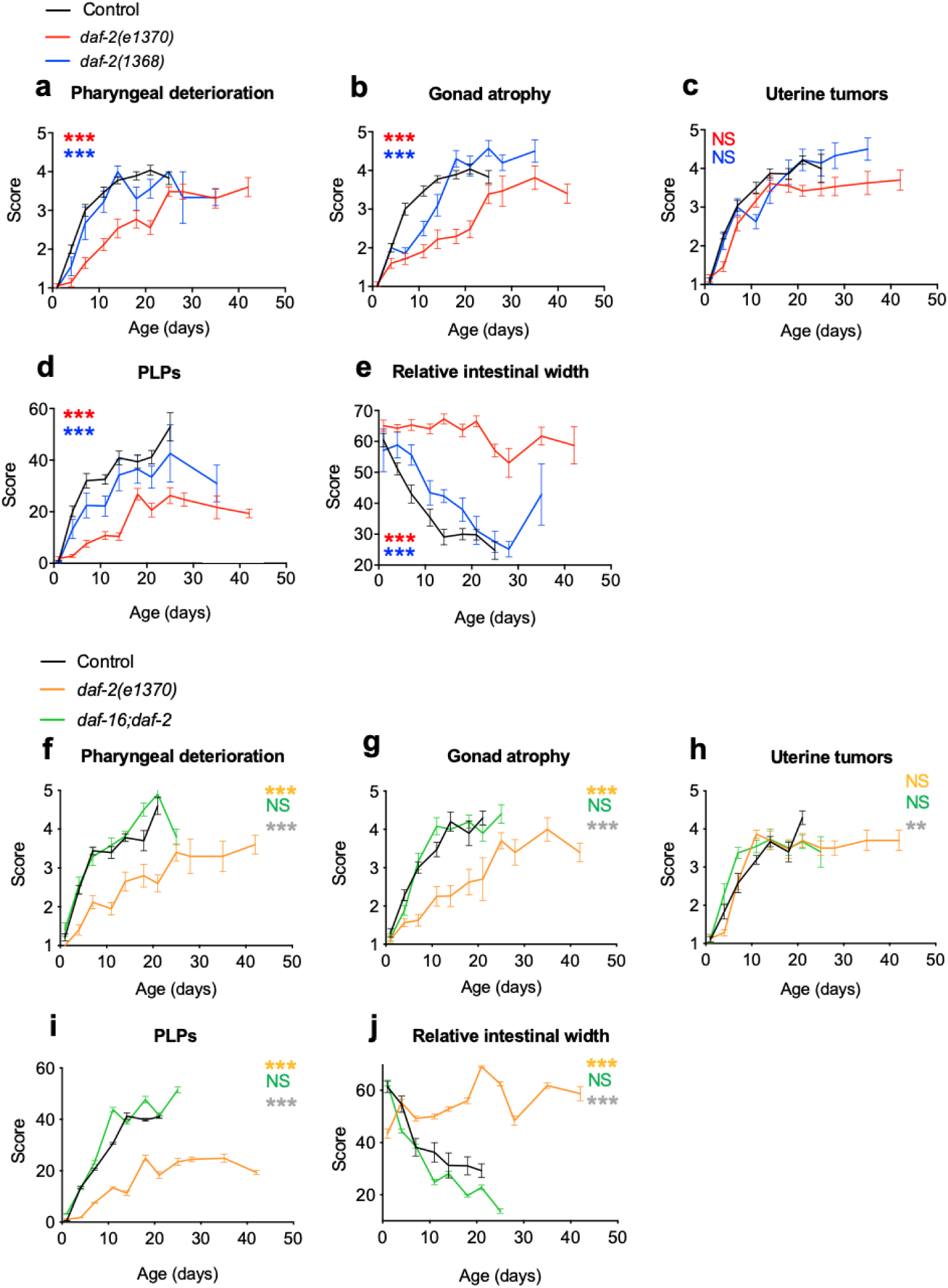
Mutation of *daf-2* differentially suppresses senescent pathology. (a-e) *daf-2(e1370)* suppresses pathology development more strongly than *daf-2(e1368)*. (a) Pharyngeal deterioration, (b) gonad disintegration, (c) uterine tumors, (d) PLPs (yolk pools), (e) gut atrophy. (f-i) Pathology suppression by *daf-2(e1370)* is dependent on DAF-16. (f) Pharyngeal deterioration, (g) gonad disintegration, (h) uterine tumors, (i) PLPs, (j) gut atrophy. 2 trials (n=10). Two-way ANOVA (Tukey correction). Stars denote differences in pathology progression via ANCOVA (Tukey correction). **** *p* < 0.00001; *** *p* < 0.0001; ** *p* < 0.001; * *p* < 0.01.

**Supplementary Figure 4.**
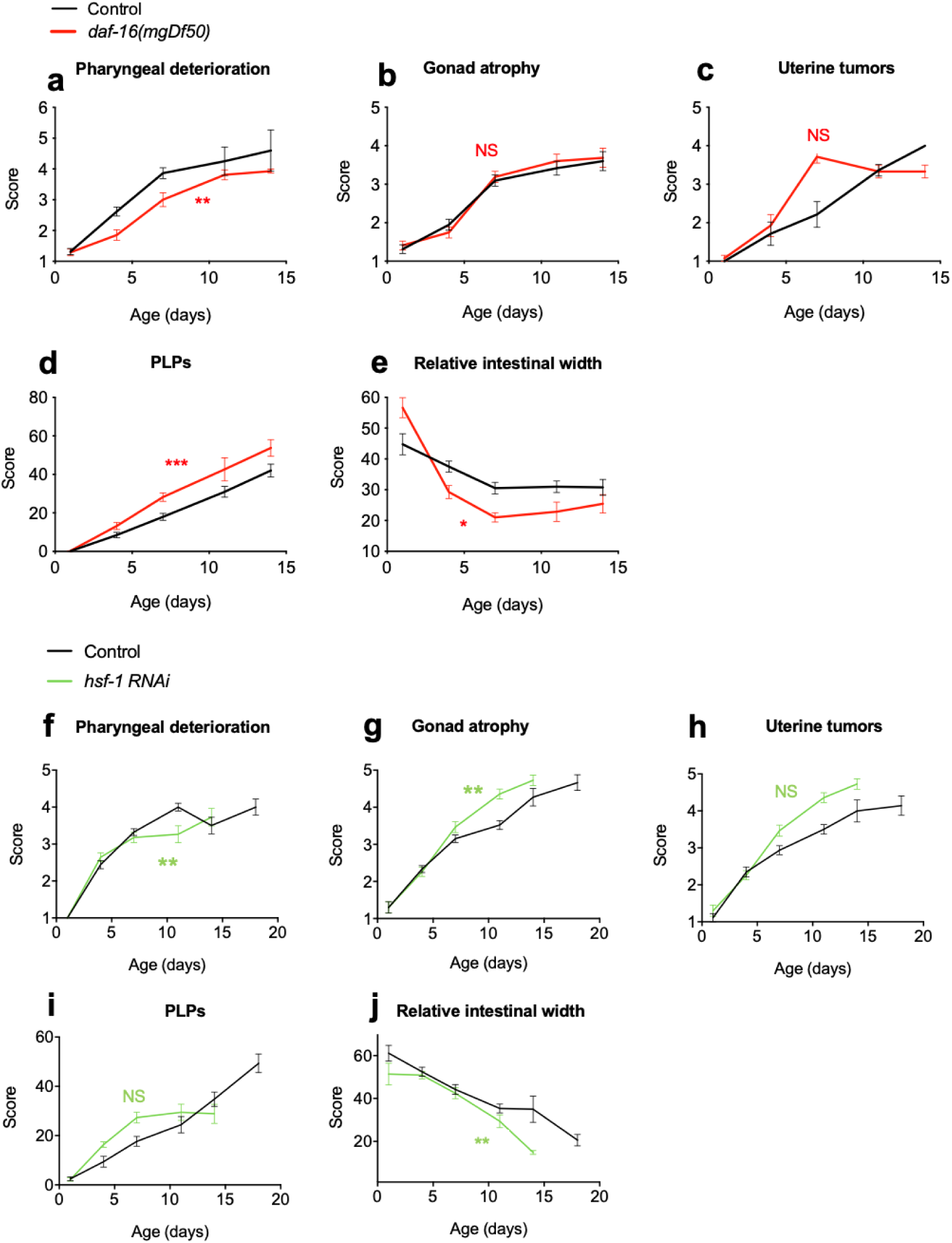
Effects on pathology development of tissue-specific *daf-16* rescue and *hsf-1* inhibition. (a-e) *daf-16(mgDf50)* accelerates some but not all pathologies. (a) Pharyngeal deterioration (b) Gonad atrophy. (c) Uterine tumors. (d) PLPs. (e) Relative intestinal width. (f-j) *hsf-1* RNAi accelerates some but not all pathologies. (f) Pharyngeal deterioration. (g) Uterine tumors. (h) PLPs. (i) Relative intestinal width. 2 trials (n=10). Two-way ANOVA (Bonferroni correction), and stars show differences in pathology progression via ANCOVA (Tukey correction). * *p*<0.01; ** *p*<0.001; *** *p*<0.0001; **** *p*<0.00001.

**Supplementary Figure 5.**
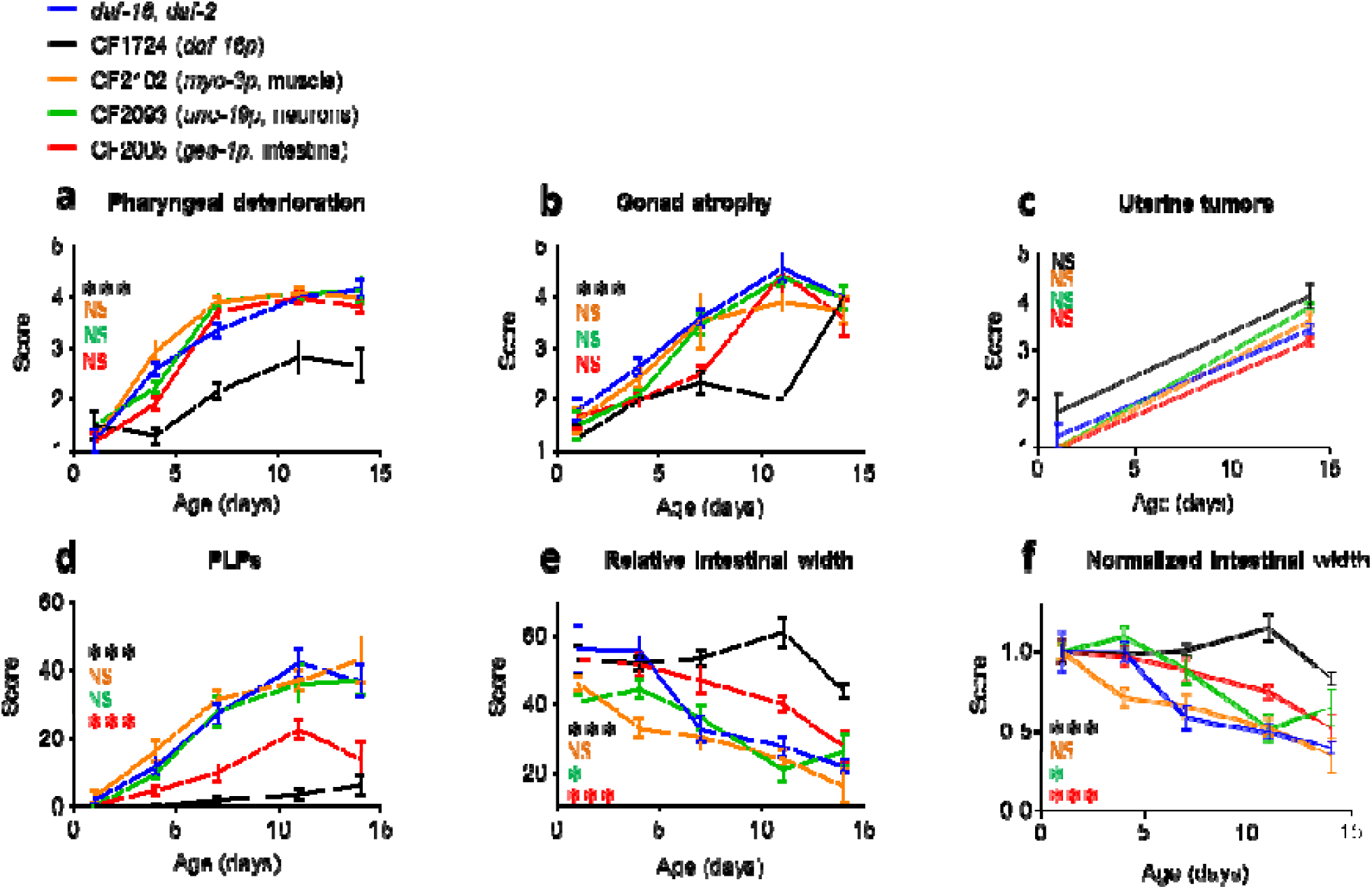
Effects of tissue-specific rescue of *daf-16* on senescent pathology in *daf-16; daf-2(e1370)* mutants. *myo-3p::* muscle*; ges-1p::* intestine; and *unc-119p::* neurons. (a) Pharyngeal deterioration, (b) gonad atrophy, (c) uterine tumors, (d) PLPs, (e) gut atrophy. (f) Relative intestinal width normalised to intestinal score on day 1 per condition. Two-way ANOVA (Tukey correction), and stars show differences in pathology progression via ANCOVA (Tukey correction). * *p*<0.01; ** *p*<0.001; *** *p*<0.0001; **** *p*<0.00001.

**Supplementary Figure 6.**
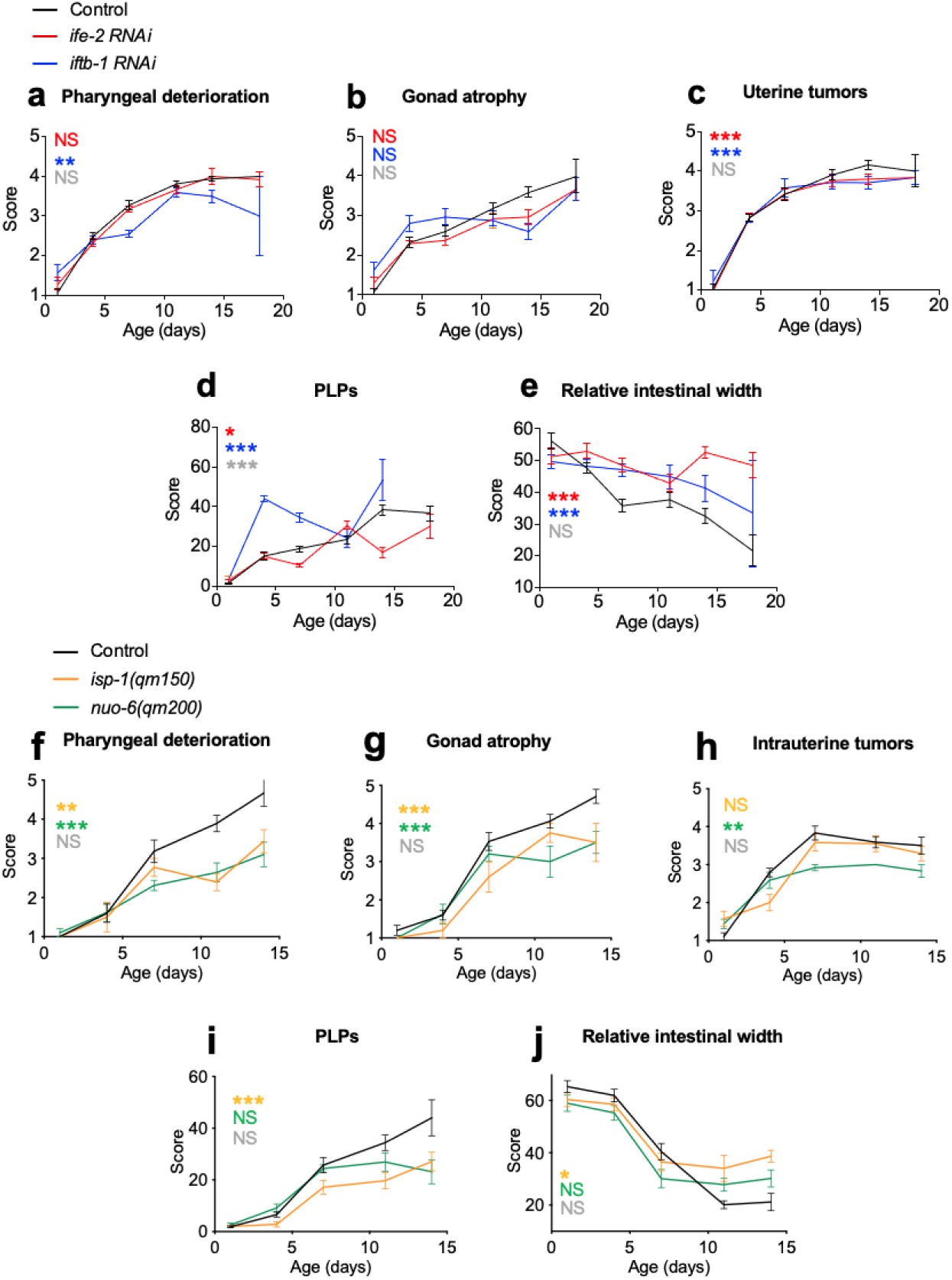
Evidence that protein synthesis and mitochondrial function promote pathology development. (a-e) *ife-2* and *iftb-1* RNAi inhibits gut atrophy but not most other pathologies. (a) Pharyngeal deterioration, (b) gonad disintegration, (c) uterine tumors, (d) PLPs (yolk pools), (e) gut atrophy. (f-j) *isp-1(qm150)* and *nuo-6(qm200)* inhibit multiple age-related pathologies. (f) Pharyngeal deterioration, (g) uterine tumors, (h) PLPs (yolk pools), (i) gut atrophy. 2 trials (*n*=10). Two-way ANOVA (Tukey correction). Stars show differences in pathology progression via ANCOVA (Tukey correction). The color of the stars represents treatment color vs control. Grey stars: between treatment comparison. * *p* < 0.01; ** *p* < 0.001; *** *p* < 0.0001; **** *p* < 0.00001.

**Supplementary Figure 7.**
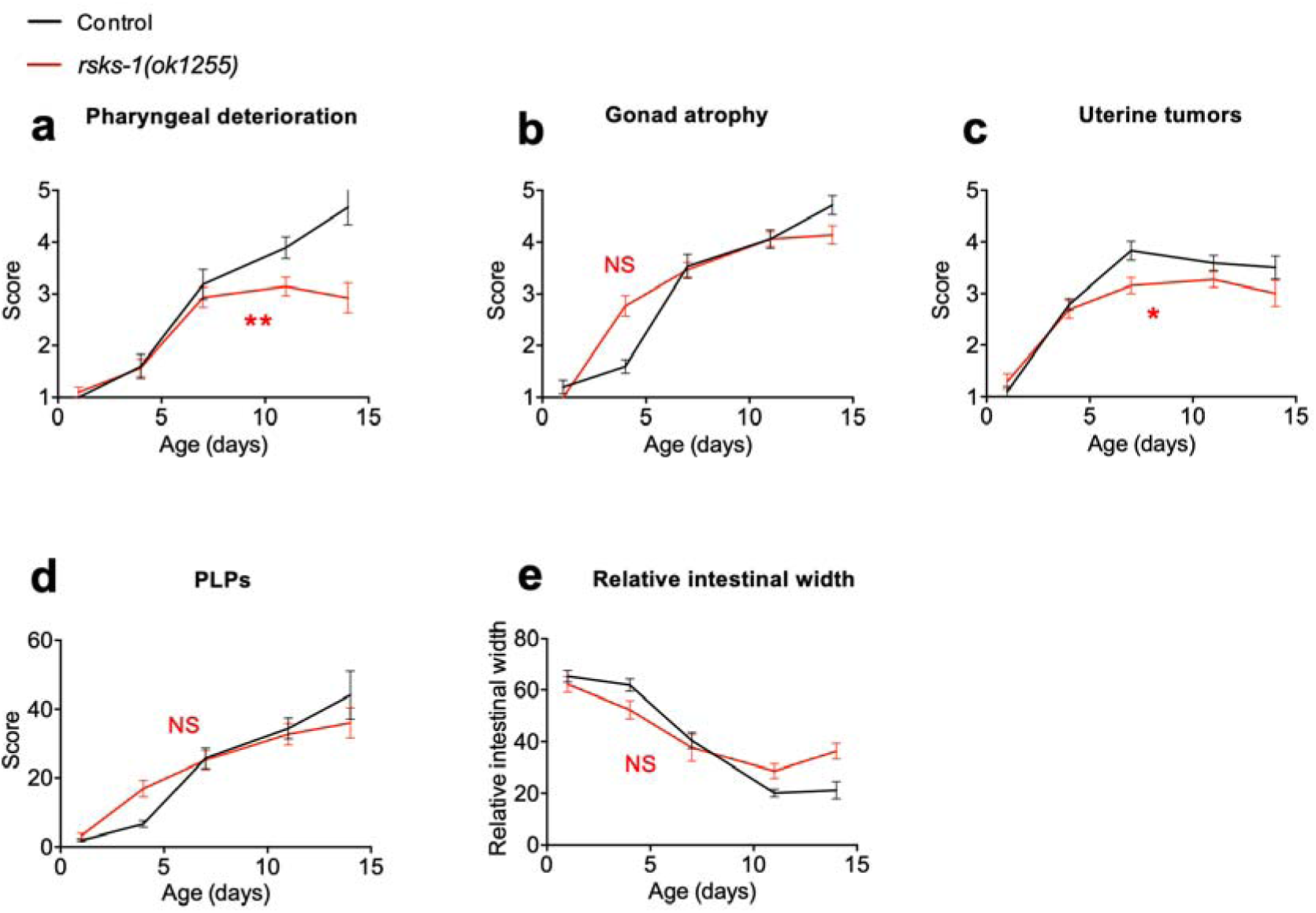
Ribosomal protein S6 kinase has small effects on pathology development. (a-e) *rsks-1(ok1255)* suppresses pharyngeal decline and tumor formation but not other pathologies. (a) Pharyngeal deterioration. (b) Gonad atrophy. (c) Uterine tumors. (d) PLPs. (e) Relative intestinal width. Two-way ANOVA (Tukey correction), and stars show differences in pathology progression via ANCOVA (Tukey correction). * *p*<0.01; ** *p*<0.001; *** *p*<0.0001; **** *p*<0.00001.

**Supplementary Figure 8.**
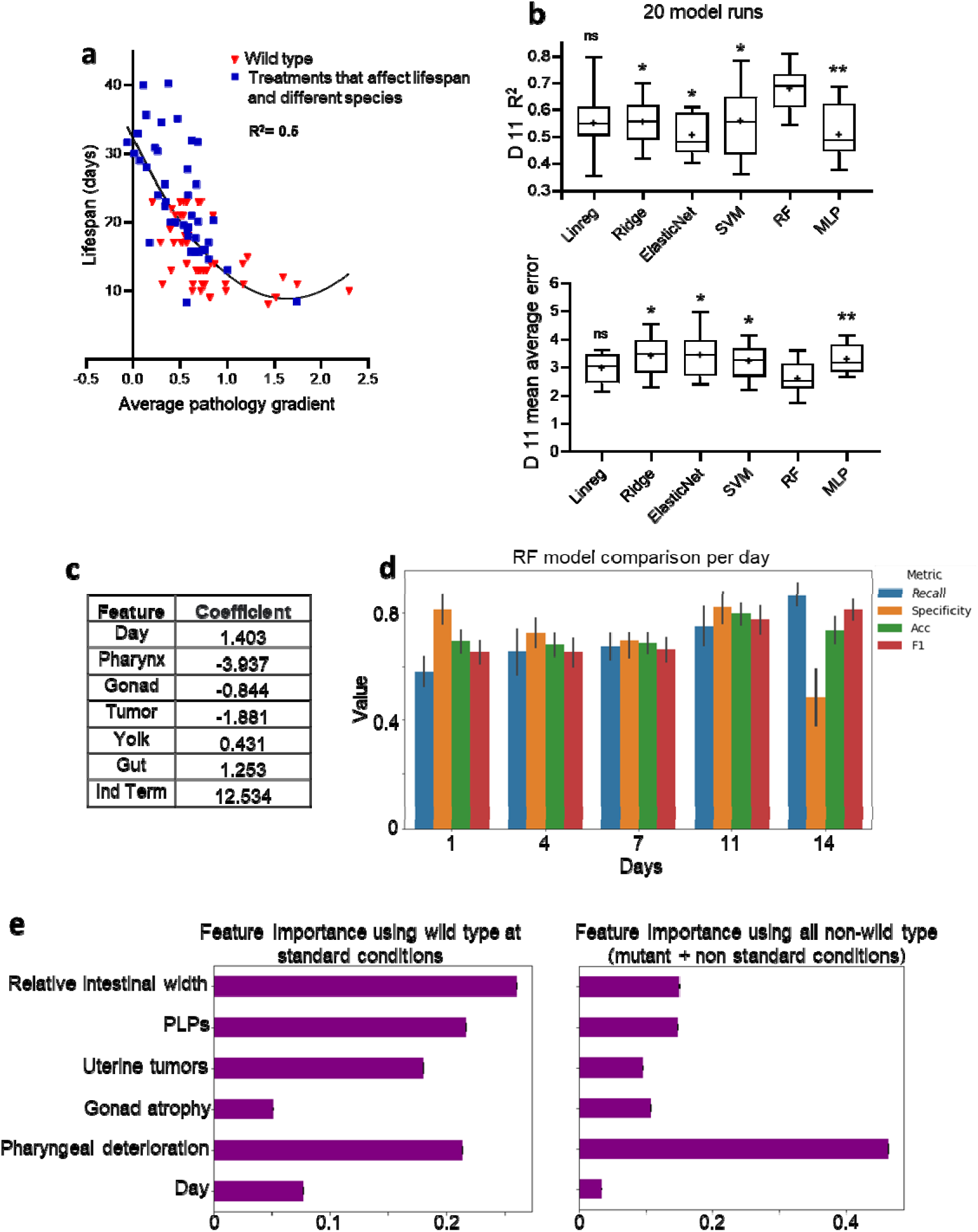
Senescent pathology can be used to predict lifespan. (a) Broad correlation between pathology and lifespan. Pathology progression through time is converted into a gradient and then transformed into Z-scores (which describe a value’s relationship to the mean of a group of values) for standardisation. This allows comparison between different pathologies by normalising levels of a given pathology to the average level of that pathology in the group (represented by a Z-score of 0, with a Z-score of +2 representing the maximum pathology severity in the group and -2 the healthiest animals). Only intestine scores were normalised to the respective mean intestine score on day 1 prior to being converted to a gradient, so the normalised intestinal pathology scores represent the change in percentage of intestinal volume and this accounts for differences in terms of the ratio of intestinal width to whole body width between different treatments and species. The average of all 5 pathology Z-scores is shown (x-axis) plotted against real observed lifespan (y-axis). (b) R^2^ and mean average error (days) with different ML models (linear regression, ElasticNet regression, Support Vector Machine [SVM], random forest [RF], multilayer perceptron [MLP]) and pathology progression accounted until day 11 of adulthood. pval comparing RF model to other models. *** *p*<0.0001; **** *p*<0.00001. (c) Coefficients generated by the linear regression model created by accounting for pathology progression to day 11(c.f. Figure 6a). (d) Specificity and sensitivity analysis of our model derived from predicting long lived (> 18 days survival) vs short lived (<18 days survival animals). Recall =Sensitivity; F1 = F1 score; Acc = Accuracy. (e)) Feature importance: RF model Mean Decrease in Impurity for individual pathologies and day of scoring. Right: mutant animals and animals at non-standard culture conditions; Left: wild type at standard culture conditions.

**Supplementary Table 1.** Pathology scoring system. c.f. Supplementary Figure 1.

| **Score** | **Pharyngeal deterioration** | **Gonad atrophy** | **Uterine tumours** |
| --- | --- | --- | --- |
| **1** | Healthy pharynx. Borders are smooth, radial muscle lines are visible and intact. | Healthy gonad. Gonad arms are intact. Edges are smooth. Germine cells are surrounded by cytoplasm. Gonad arms fill up the body cavity and are mostly touching each other. | Healthy uterus, no tumours. |
| **2** | Borders of the pharynx are uneven. Radial muscle lines are starting to fade. Small vacuoles/cavities appear. | Thinned gonad. Starting to show deterioration. Diameter of gonad arm is decreased and are touching less than in a healthy gonad. Edges of distal arm are starting to shrivel. | Abnormal uterus; possible beginning of tumour |
| **3** | Aged pharynx. One of the following ageing features: bacterial plug present, terminal bulb starting to swell or cavities in bulb. | Aged gonad. Gonad even thinner. Further decrease of diameter and further shrivelling. | Small tumours: $\text{Area}/(\text{diameter})^2 < 0.9$ |
| **4** | Very aged. Two or more of the ageing features above. | Very aged. Fragmented gonad. Disruption/displacement/ twisting of gonad | Large tumours but do not fill entire cavity; $A/d^2 > 0.9$ |
| **5** | Posterior pharyngeal bulb completely swollen or disintegrated. | Gonad not recognisable. | Massive tumours; fills the entire diameter of body cavity |

**Supplementary Table 2.** All lifespan data sources and pathology data sources collated and used in ML analysis. C.f. Figure 3.

| Treatment, species, genotype | Pathology data source | Lifespan data source |
| --- | --- | --- |
| Culture at 15°C | This study | 36 |
| Culture at 25°C | This study | 36 |
| Culture on solid axenic medium | This study | 5 |
| Mated hermaphrodites | This study | 37 |
| Male | This study | 12 |
| *C. elegans* | 38 | 38 |
| *C. inopinata* | 38 | 38 |
| *C. briggsae* | 38 | 38 |
| *C. nigoni* | 38 | 38 |
| *C. remanei* | 38 | 38 |
| *C. tropicalis* | 38 | 38 |
| *C. wallacei* | 38 | 38 |
| *C. remanei* | This study | This study |
| *Pristionchus exspectatus* | 38 | 38 |
| *Pristionchus pacificus* | 38 | 38 |
| Laser ablation z1-4 *C. elegans* | 38 | 38 |
| Laser ablation z23 *C. briggsae* | 38 | 38 |
| Laser ablation z23 *C. elegans* | 38 | 38 |
| Laser ablation z23 *C. tropicalis* | 38 | 38 |
| Laser ablation z23 *P. pacificus* | 38 | 38 |
| *glp-1(bn4)* | 38 | 32 |
| *glp-4(bn2)* | 38 | 39 |
| *daf-2(e1370)* | This study | 40 |
| *daf-2(e1368)* | This study | 11 |
| *daf-16; daf-2* | This study | 40 |
| *daf-16(mgDf50)* | This study | 40 |
| *Pdaf-16::daf-16; daf-16* | This study | 20 |
| *Pges-1::daf-16; daf-16* | This study | 20 |
| *Pmyo-3::daf-16; daf-16* | This study | 20 |
| *Punc-119::daf-16; daf-16* | This study | 20 |
| *mab-3* | 11 | 11 |
| *mab-3 males* | 11 | 11 |
| *him-5* | This study | 11 |
| *him-5 males* | This study | 11 |
| *fem-3* | 38 | 38 |
| *fog-2* | 38 | 38 |
| *rps-15* RNAi | This study | 25 |
| *iftb-1* RNAi | This study | 25 |
| *rsks-1(ok1255)* | This study | 28 |
| *isp-1* | This study | 31 |
| *nuo-6* | This study | 31 |
| *hsf-1* RNAi | This study | 41 |
| *ife-2* RNAi | This study | 27 |
| *iftb-1* RNAi | This study | 25 |
| *vit-5* RNAi | 14 | 14 |
| *vit-6* RNAi | 14 | 14 |
| *vit-5, vit-6* RNAi | 14 | 14 |

